# Deletion of *EP4* in *S100a4*-lineage cells reduces scar tissue formation during early but not later stages of tendon healing

**DOI:** 10.1101/132860

**Authors:** Jessica E. Ackerman, Katherine T. Best, Regis J. O’Keefe, Alayna E. Loiselle

## Abstract

Tendon injuries heal via scar tissue rather than regeneration. This healing response forms adhesions between the flexor tendons in the hand and surrounding tissues, resulting in impaired range of motion and hand function. Mechanistically, inflammation has been strongly linked to adhesion formation, and Prostaglandin E2 (PGE2) is associated with both adhesion formation and tendinopathy. In the present study we tested the hypothesis that deletion of the PGE2 receptor *EP4* in *S100a4*-lineage cells would decrease adhesion formation. S100a4-Cre; *EP4*^flox/flox^ (EP4cKO^S100a4^) repairs healed with improved gliding function at day 14, followed by impaired gliding at day 28, relative to wild type. Interestingly, EP4cKO^S100a4^ resulted in only transient deletion of *EP4,* suggesting up-regulation of *EP4* in an alternative cell population in these mice. Loss of *EP4* in *Scleraxis-*lineage cells did not alter gliding function, suggesting that *Scx*-lineage cells are not the predominant *EP4* expressing population. In contrast, a dramatic increase in α-SMA+, EP4+ double-positive cells were observed in EP4cKO^S100a4^ suggesting that EP4cKO^S100a4^ repairs heal with increased infiltration of EP4 expressing α-SMA myofibroblasts, identifying a potential mechanism of late up-regulation of *EP4* and impaired gliding function in EP4cKO^S100a4^ tendon repairs.

## Introduction

Tendons are dense connective tissues composed primarily of a highly organized type I collagen extracellular matrix (ECM). The hierarchical structure of the collagen ECM allows tendons to transmit large forces between muscle and bone, facilitating movement of nearly the entire body. Acute traumatic injuries to the tendon are quite common due in part to the superficial anatomic location of many tendons. Following injury, tendons generally heal with a scar tissue response rather than regeneration of native tendon. This fibrotic healing response is particularly problematic in the hands, as excursion of the flexor tendons within the synovial sheath is restricted by both increased tissue bulk and the formation of adhesions between the tendon and surrounding tissues. Currently, treatment to restore digit range of motion (ROM) after injury is limited to surgical ‘tenolysis’ procedures ^1^, however this intervention provides only a temporary return to function. Thus, a better understanding of the fundamental cellular and molecular components of fibrotic healing are needed in order to inform development of future therapeutic interventions to improve tendon healing.

Tendon healing is composed of three over-lapping phases of healing including inflammation, proliferative/ matrix deposition, and the remodeling phase. Several studies have demonstrated that anti-inflammatory treatment, particularly Cox-2 inhibition, effectively reduces adhesion formation ^2,3^, however, mechanical properties can also be compromised with this approach ^2^. Therefore, recent studies have focused on downstream signaling of Cox-2 as potential therapeutic targets. Cox-2 catalyzes the conversion of Arachidonic Acid to Prostaglandins, including PGE2, which is associated with tendinopathy and involved in tendon repair ^4–8^. Moreover, EP4 has been identified as the specific receptor through which degenerative tendinopathic changes of PGE2 are mediated ^9^. We have previously demonstrated that systemic EP4 antagonism increased adhesion formation, relative to vehicle treated repairs ^10^, however the specific function of EP4 is dependent on cell type ^11,12^. Therefore, in the present study we have focused on the effects of cell-type specific EP4 deletion on tendon healing.

In the present study we utilized *S100a4-Cre; EP4*^flox/flox^ (EP4cKO^S100a4^) mice to test the hypothesis that conditional deletion of *EP4* in *S100a4-lineage* cells would reduce adhesion formation while maintaining mechanical properties, relative to wildtype (WT) littermates. S100a4 (S100 Calcium Binding Protein A4) is a member of the EF-hand Ca2+-binding proteins, and is associated with fibrosis in several tissues ^13–16^. Given the fibrotic nature of scar-mediated tendon healing, we determined the feasibility of using S100a4-Cre as an efficient means to target the tendon during healing, and demonstrate recombination in both the un-injured tendon, and at the tendon repair site. We show that flexor tendon repairs from EP4cKO^S100a4^ have only a transient, early inhibition of *EP4* expression, concomitant with early reductions in adhesion formation. However, during later healing *EP4* expression is increased in EP4cKO^S100a4^ repairs relative to WT, suggesting an alternative cell population up-regulates *EP4* in these mice. We then show that loss of *EP4* in *Scleraxis-*lineage cells does not alter adhesion formation, while EP4cKO^S100a4^ repairs have an increase in the myofibroblast marker α-SMA, suggesting a potential mechanism of increased adhesion formation that occurs during later healing in EP4cKO^S100a4^ mice.

## Results

### S100a4-lineage cells are found in both native tendon and the granulation tissue

To determine the expression pattern of S100a4-lineage cells using S100a4-Cre, and thus the potential localization of EP4 conditional deletion, the *ROSA*^nT/nG^ dual reporter was used. GFP-expressing (S100a4-Cre^+^) cells were observed in 40.5% of resident tenocytes in the absence of injury (Fig. 1A, D). At day 7 post-surgery abundant GFP^+^, S100a4-Cre^+^ cells were observed throughout the native tendon and within the highly cellular bridging granulation tissue (Fig. 1B & B’, Supplemental Figure 1), with 79.9% of cells in the scar tissue being nGFP^+^ (S100a4^+^) (p<0.001 vs. un-injured)(Fig. 1D). At day 14 post-repair a few GFP^+^, S100a4-Cre^+^ cells were observed in the ends of the native tendon, however, abundant GFP^+^, S100a4-Cre^+^ cells were present in the granulation tissue that bridges the injury site (Fig. 1C & 1C’), with 74.7% of cells in the scar tissue being nGFP^+^ (S100a4^+^) (p<0.001 vs un-injured)(Fig. 1D).

**Figure 1.**
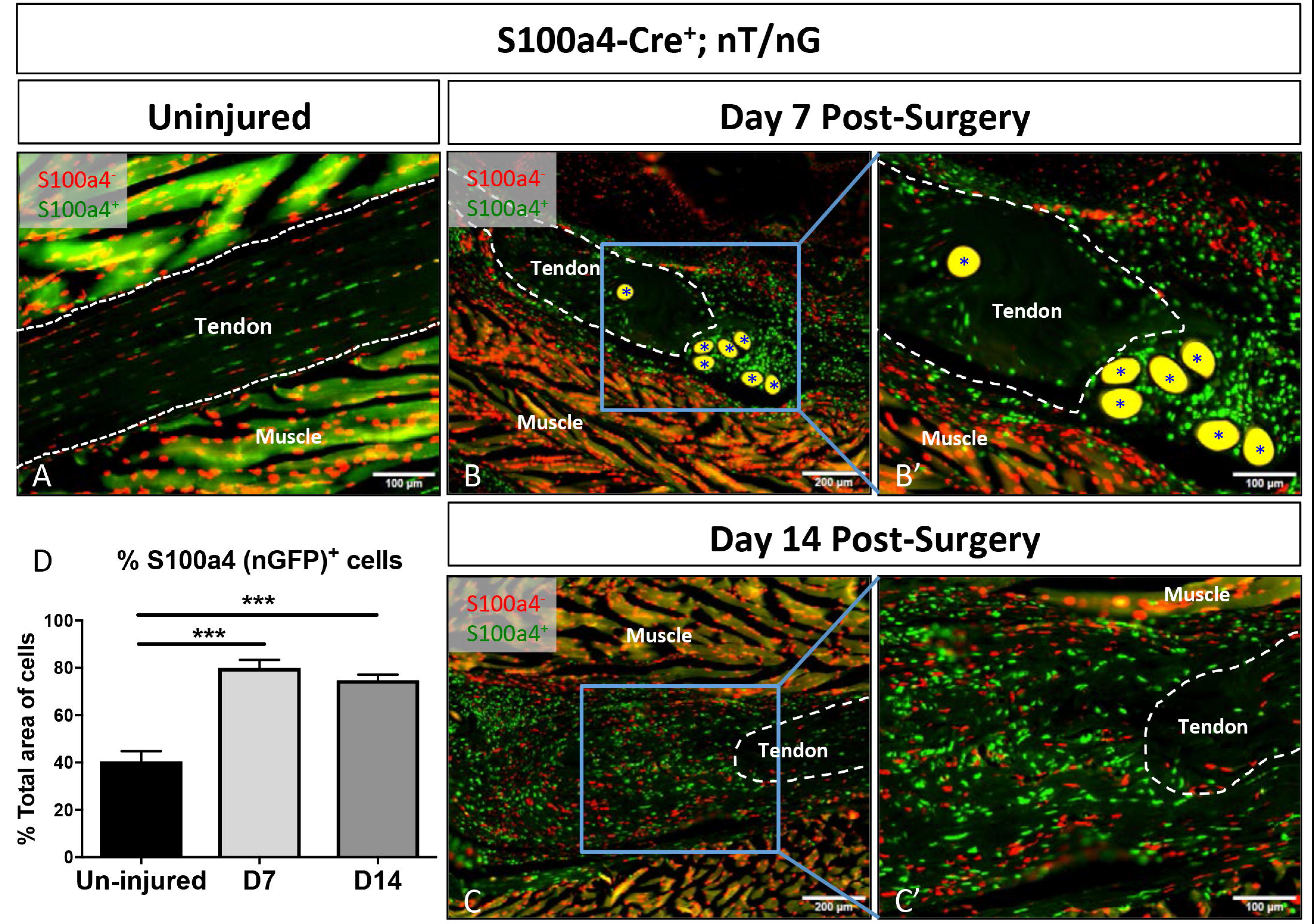
S100a4-Cre effectively targets tendon. To demonstrate that *S100a4*-Cre effectively targets recombination in the tendon we examined healing in the double reporter nT/nG with *S100a4*-Cre^+^ cells expressing GFP, and *S100a4*-Cre^-^ cells expressing Td Tomato Red. *S100a4*-Cre results in recombination in both the (A) un-injured and (B & C) repaired tendon at days 7 (B & B’) and 14 (C & C’) post-surgery. Tendon tissue is outlined in white, (*) indicate auto-fluorescent sutures at the repair site. Scale bars (A, B’, C’) represent 100 microns, and (B & C) 200 microns. (D) Quantification of the proportion of nGFP^+^ (*S100a4*-Cre^+^) cells in the un-injured tendon and at 7 and 14 days post-repair. (***) Indicates p<0.001 vs. un-injured.

### EP4cKO^S100a4^ Does not alter mechanical properties of un-injured tendons

To ensure that EP4cKO^S100a4^ did not impair baseline mechanical properties and gliding function of the FDL, uninjured tendons were used. No change in Max load at failure (WT: 7.33N ± 0.35; EP4cKO^S100a4^: 7.89 ± 0.67 p=0.46) (Fig. 2A), stiffness or energy to Max (data not shown) were observed between un-injured EP4cKO^S100a4^ and WT tendons. Additionally, no change in MTP flexion angle (WT: 51.98 ± 1.75; EP4cKO^S100a4^: 51.78 ± 3.95, p=0.96) (Fig. 2B) and Gliding Resistance (WT: 9.63 ± 0.95; EP4cKO^S100a4^: 11.09 ± 2.11, p=0.53) were observed between genotypes (Fig. 2C).

**Figure 2.**
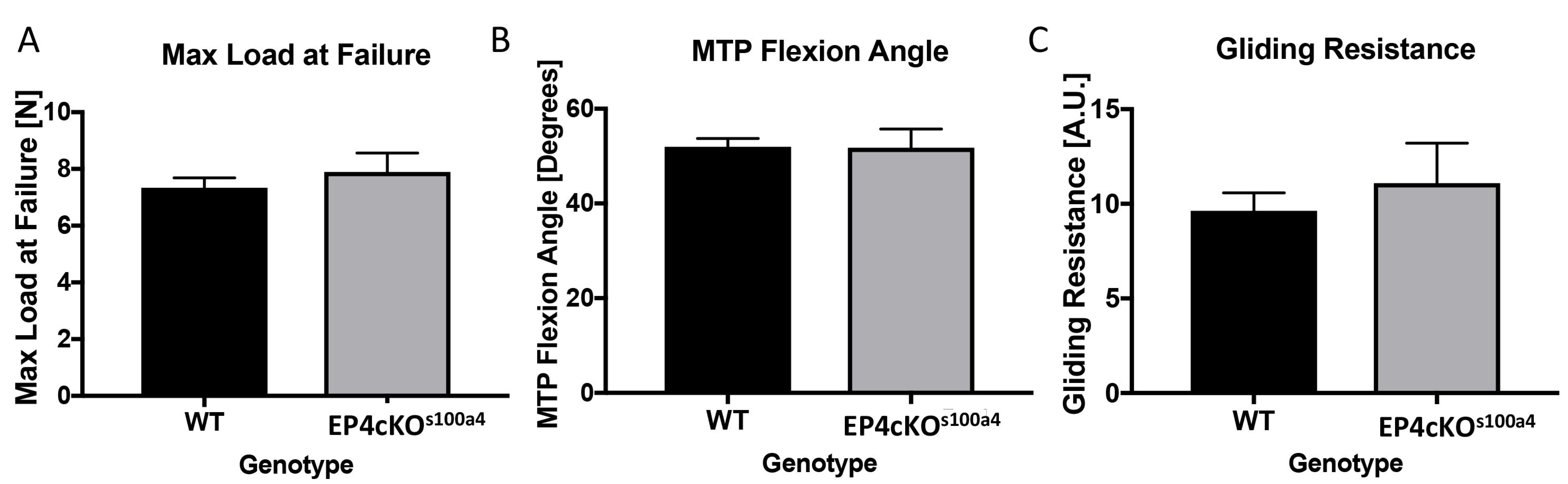
EP4cKO^S100a4^ does not alter gliding function or mechanical properties of un-injured tendons. No changes in (A) Max load at failure, (B) MTP Flexion Angle, or (C) Gliding Resistance were observed betweenun-injured tendons from WT and EP4cKO^S100a4^ flexor tendons.

### EP4 expression is decreased during early but not late healing in EP4cKO^S100a4^ tendon repairs

To determine the temporal effects of EP4cKO^S100a4^ on *EP4* expression during healing, mRNA was isolated from WT and EP4cKO^S100a4^ repairs between 7 and 28 days post-repair. Peak *EP4* expression was observed in WT repairs at day 7 post-repair, consistent with previous studies ^10^, followed by a significant reduction in *EP4* expression at 14, 21 and 28 days, relative to day 7. A significant 63.9% reduction in *EP4* expression was observed in EP4cKO^S100a4^ repairs relative to WT (p=0.0135, Fig. 3) at day 7, identifying efficient reduction in *EP4* expression at this time. In contrast, *EP4* expression was significantly increased in EP4cKO^S100a4^ repairs at days 14 and 21, relative to time-point matched WT repairs. No change in *EP4* expression was observed between WT and EP4cKO^S100a4^ repairs at day 28 (Fig. 3)

**Figure 3.**
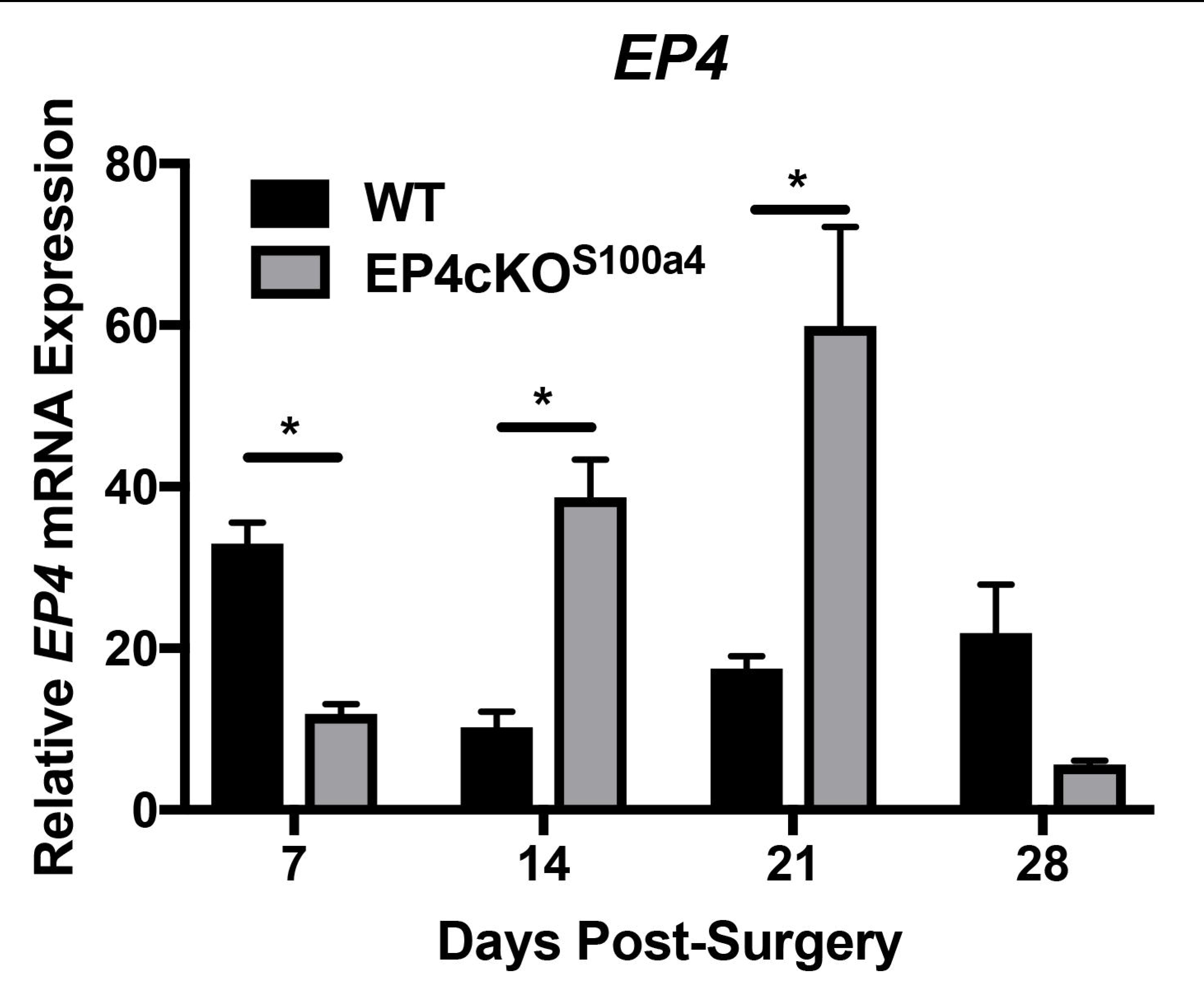
EP4cKO^S100a4^ significantly decreases *EP4* expression during early healing. mRNA was isolated from EP4cKO^S100a4^ and WT tendon repairs between 7-28 days post-surgery. Relative *EP4* expression was significantly decreased in EP4cKO^S100a4^ relative to WT at day 7, and was significantly increased on days 14 and 21 post-surgery. Expression was normalized to the internal control *β-actin.* (*)Indicates p<0.05.

### EP4cKO^S100a4^ flexor tendon repairs have an early, transient improvement in gliding function

No differences in MTP flexion angle were observed at day 10 post-repair between WT and EP4cKO^S100a4^ mice. However, by day 14 a significant 71% increase in MTP Flexion angle was observed in EP4cKO^S100a4^ repairs (WT: 14.53 ± 1.74; EP4cKO^S100a4^: 28.55 ± 5.44, p=0.03). At 21 days MTP flexion was not different between genotypes, however, at 28 days MTP Flexion was significantly increased in WT repairs relative to EP4cKO^S100a4^ repairs (p=0.03) (Fig. 4A). A significant increase in Gliding Resistance occurred in WT repairs at day 14, relative to EP4cKO^S100a4^ (WT: 65.00 ± 8.06; EP4cKO^S100a4^: 37.60 ± 7.887 p=0.034) (Fig. 4B). No changes in Gliding Resistance were observed between groups at any other time-points.

**Figure 4.**
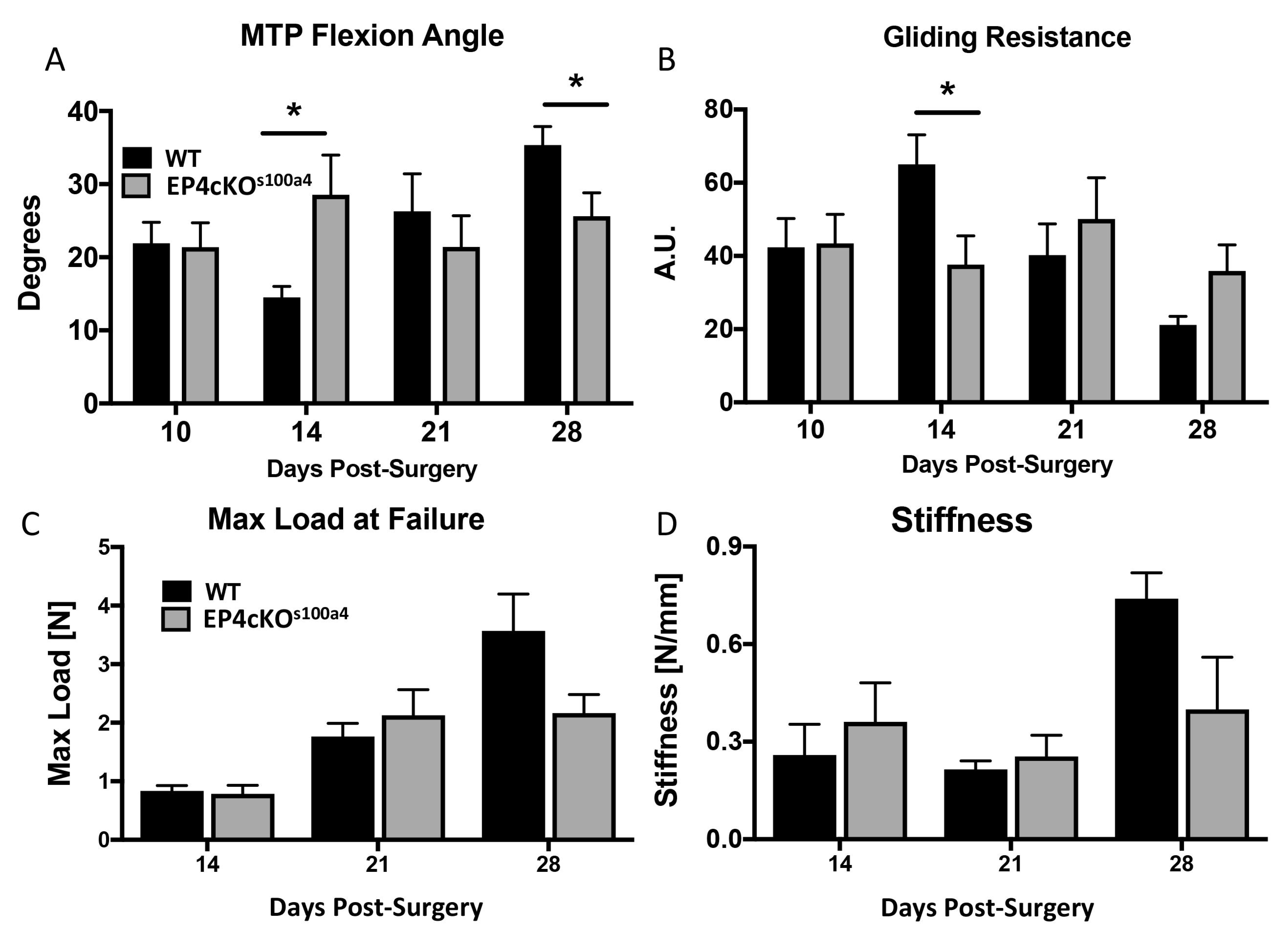
EP4cKO^S100a4^ significantly improves early gliding function without compromising mechanical properties during flexor tendon healing. (A) MTP Flexion Angle, and (B) Gliding Resistance were measured in WT and EP4cKO^S100a4^ flexor tendons between 10-28 days post-surgery. (C) Max load at failure, and (D) Stiffness were assessed between 14-28 days post-surgery. (*) Indicates p<0.05 between WT and EPcKO^S100a4^ at the same time-point.

### EP4cKO^S100a4^ does not impair mechanical properties during flexor tendon healing

Max load at failure improved progressively in both WT and EP4cKO^S100a4^ repairs from day 14 to day 28. While no differences were observed in max load at failure between genotypes at a given time point, accrual of strength was slightly retarded in EP4cKO^S100a4^ repairs, and there was a trend toward decreased max load at failure in EP4cKO^S100a4^, relative to WT at Day 28 (p=0.068). The max load at failure improved 328% in WT repairs from day 14 to Day 28 (Day 14: 0.83 ± 0.09; Day 28: 3.57 ± 0.62). In contrast, EP4cKO^S100a4^ repairs regained only 176% of max load at failure from day 14 to day 28 (Day 14: 0.78 ± 0.14; Day 28: 2.164 ± 0.31) (Fig. 4C). No differences in stiffness occurred between WT and EP4cKO^S100a4^ repairs at any time, however, there was a trend toward an increase in stiffness in WT, relative to EP4cKO^S100a4^ at day 28 (WT: 0.739 ± 0.07; EP4cKO^S100a4^: 0.399 ± 0.16, p=0.076)(Fig. 4D).

### EP4cKO^S100a4^ Repairs Heal with Decreased Expression of Collal and Col3a1

No change in *Col1a1* expression was observed between groups at day 7 post-surgery, and the typical progressive increase in *Col1a1* was observed in WT repairs between 7-21 days, followed by a return to baseline levels by day 28. In contrast, *Col1a1* was significantly decreased in EP4cKO^S100a4^ repairs at day 14 postsurgery relative to WT (WT: 7.3 ± 3.34; EP4cKO^S100a4^: 0.76 ± 0.22, p=0.03), followed by a non-significant decrease at day 21 (WT: 10.36 ± 5.11; EP4cKO^S100a4^: 5.20 ± 1.52, p=0.1) (Figure 5A). Additionally, *Col3a1* increased transiently in WT repairs from day 7-14, followed by a gradual decline back to baseline levels by day 28. *Col3a1* expression was significantly decreased in EP4cKO^S100a4^ repairs at day 14 relative to WT (WT: 5.8 ± 1.27; EP4cKO^S100a4^: 1.37 ± 0.94, p=0.04). No changes in *Col3a1* expression were observed between groups at 21 or 28 days post-surgery (Figure 5B). These data resulted in an elevated *Col1a1/Col3a1* ratio at 7 and 21 days post-surgery in WT repairs relative to EP4cKO^S100a4^.

**Figure 5.**
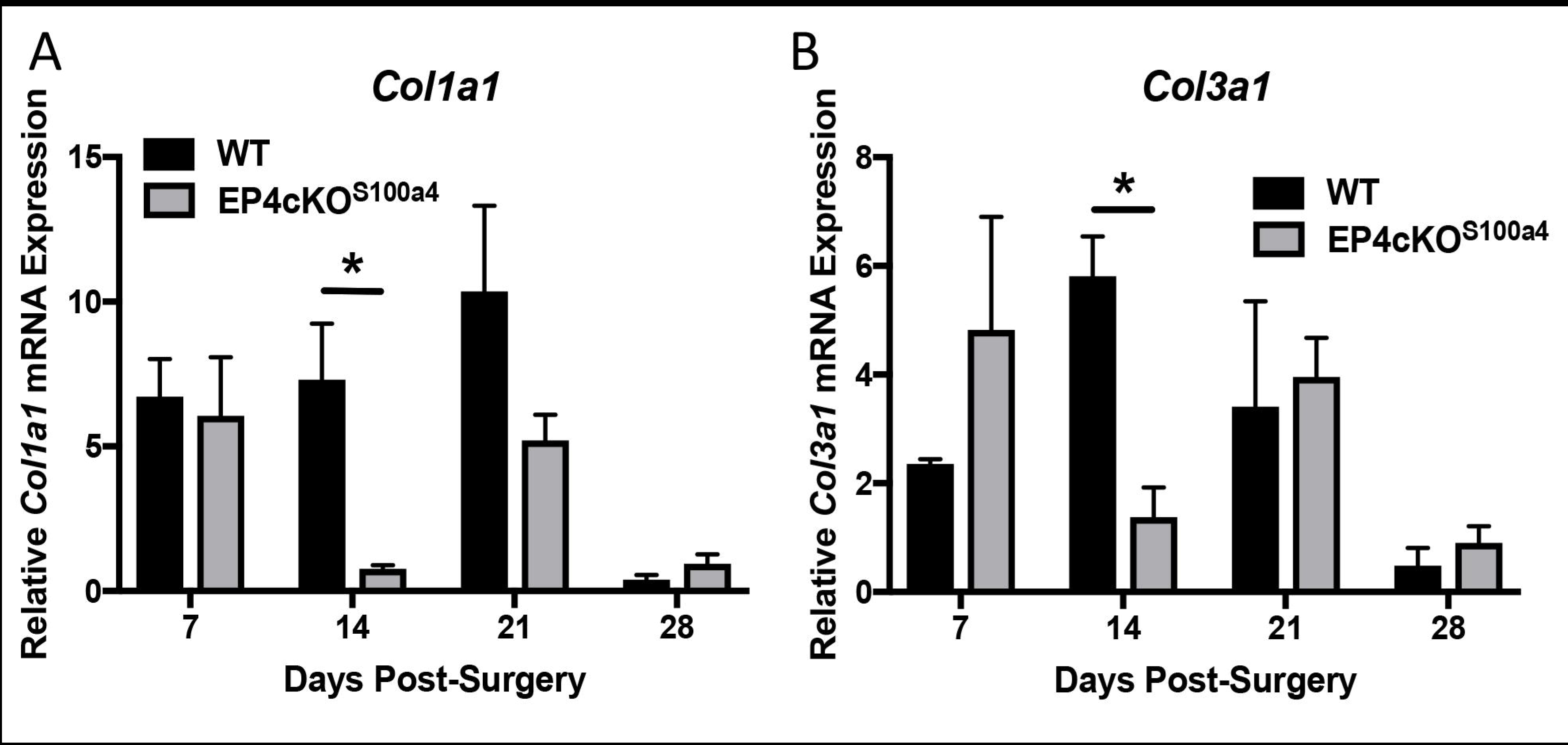
EP4cKO^S100a4^ repairs heal with decreased expression of *Collal* and *Col3a1.* mRNA was isolated from EP4cKO^S100a4^ and WT mice between 7-28 days post-surgery. A significant decrease in both (A) *Col1a1,* and (B) *Col3a1* expression was observed at day 14 post-surgery in EP4cKO^S100a4^ relative to WT. Expression was normalized to the internal control *β-actin.* (*) Indicates p<0.05.

### Loss of EP4 in Scleraxis-lineage cells does not alter flexor tendon healing

The substantial increase in *EP4* expression in EP4cKO^S100a4^ repairs suggests a cell population separate from *S100a4*-lineage expresses *EP4* during the later stages of healing. As a potential alternative population of *EP4* expressing cells, we examined the effects of *EP4* deletion in *Scleraxis*-lineage cells.

Tracing of *Scx*-lineage cells labels almost all resident tenocytes in the un-injured flexor tendon (green cells, Fig. 6A). Following injury there is an expansion of the *Scx*-lineage cells, with localization in both the native tendon and reactive epitenon, with very few *Scx*-lineage cells in the bridging scar tissue (Fig. 6B, Supplemental Figure 2).

**Figure 6.**
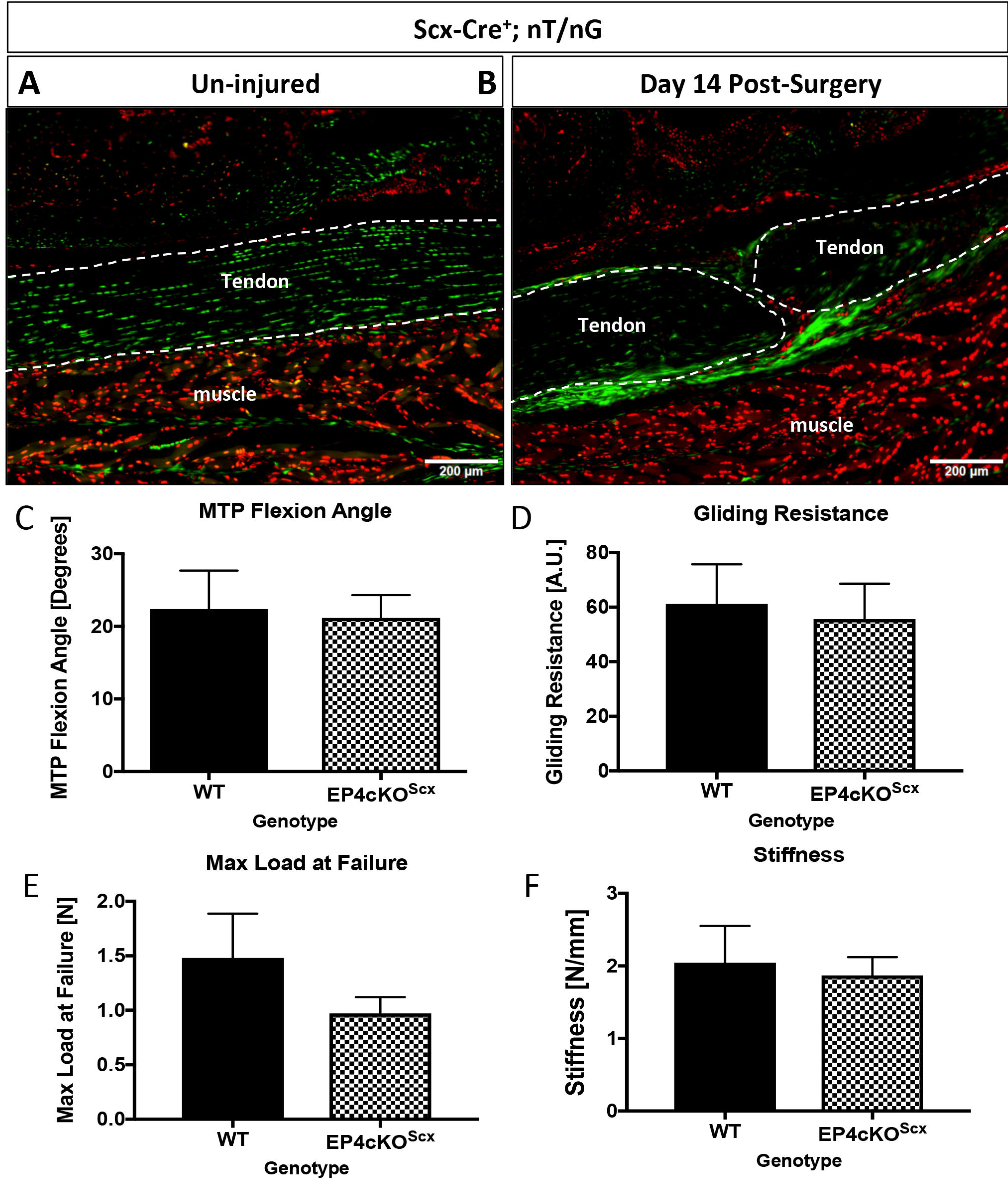
EP4cKO^Scx^ does not alter tendon healing. (A & B) Tracing of *Scx*-lineage cells using *Scx*-Cre; nT/nG demonstrates that *Scx*-lineage cells are present in the un-injured tendon (A), and the repaired tendon at day 14 post-surgery (B). Scale bar represents 100 microns. (C-F) Repaired tendons from EP4cKO^Scx^ and WT mice were harvested at day 14 post-surgery. No changes in (C) MTP Flexion Angle, (D) Gliding Resistance, (E) Max load at failure, (F) Stiffness were observed between WT repairs and EP4cKO^Scx^ repairs.

At day 14 post-repair no change in MTP flexion angle was observed between WT and EP4cKO^Scx^ (p=0.83, Fig.6C). Consistent with this, no change in Gliding Resistance was observed between genotypes (Fig. 6D). Additionally, Max load at failure (WT: 1.48N ± 0.4; EP4cKO^Scx^: 0.97N ± 0.15, p=0.17)(Fig. 6E), and Stiffness (p=0.75)(Fig. 6F) were not altered by conditional deletion of *EP4* in *Scx*-lineage cells.

### α-SMA myofibroblasts are increased in EP4cKO^S100a4^ repairs during later healing

Given that adhesion formation increases during later healing in EP4cKO^S100a4^ repairs we examined changes in pro-fibrotic, matrix producing myofibroblasts ^17^ as a potential mechanism of increased scar formation. α-SMA expressing cells were observed in both the native tendon and scar tissue of WT repairs at day 21 (Fig. 7A, Supplemental Figures 3 & 4), and no difference in α-SMA^+^ cell area was observed between WT and EP4cKOS100a4 repairs (Fig. 7C). However, abundant co-localization of α-SMA and EP4 were observed in EP4cKO^S100a4^ repairs at day 21 (yellow arrows, Fig. 7B, Supplemental Figure 4), particularly in the scar tissue, with 9.5% of the cell area positive for both α-SMA and EP4 (Fig. 7E). In contrast, very few double positive (α-SMA^+^, EP4^+^) cells were observed in WT repairs (0.61%)(Fig. 7E), consistent with a decrease in EP4^+^ cells at this time-point in WT (4.5%) relative to EP4cKO^S100a4^ (29.8%, p=0.0012)(Fig. 7D).

**Figure 7.**
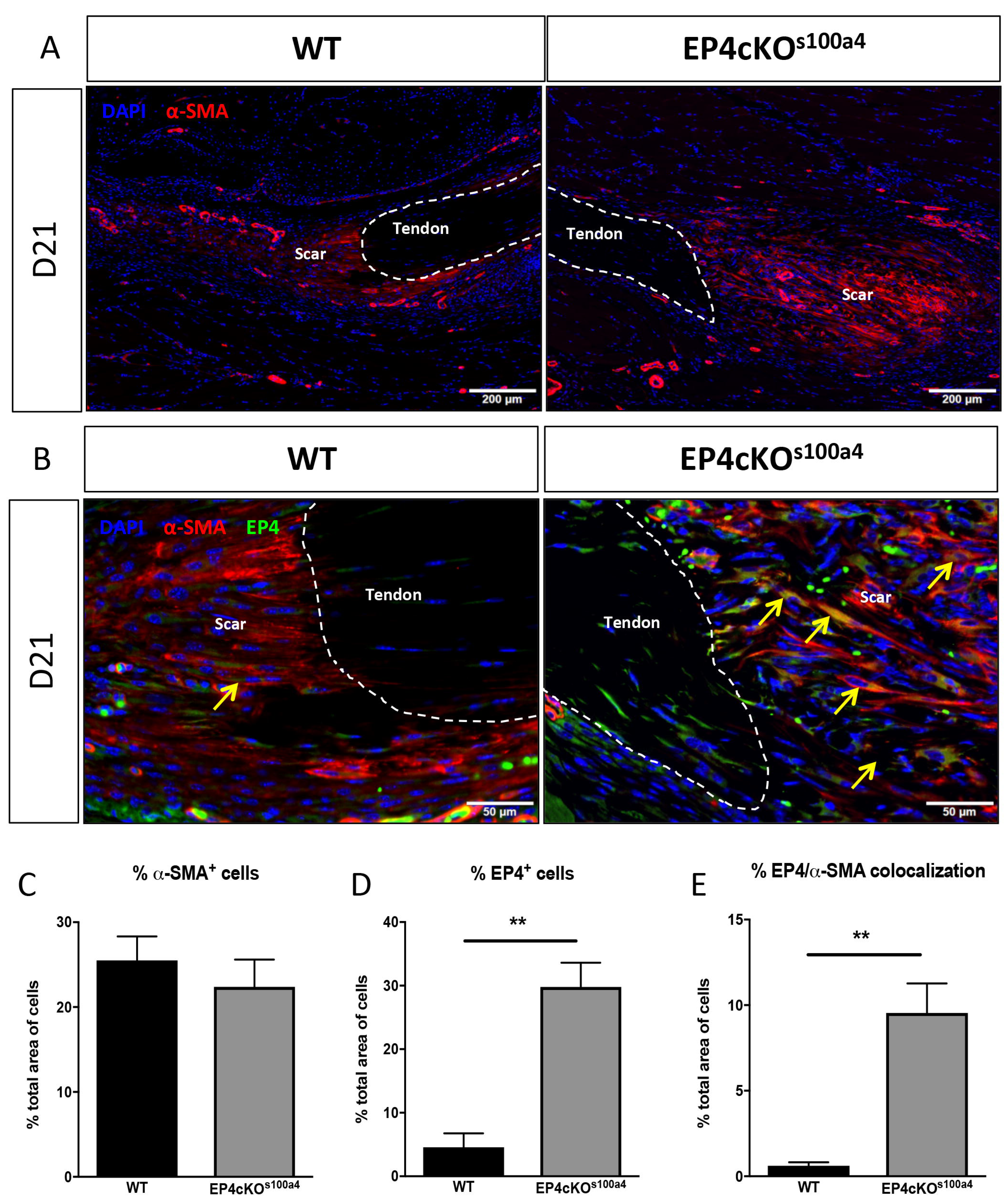
EP4-expressing α-SMA myofibroblasts are increased in EP4cKO^S100a4^ repairs during later healing. (A) Abundant expression of α-SMA^+^ was observed in EP4cKO^S100a4^ and WT repairs on day 21 postsurgery. α-SMA^+^ cells are identified by red fluorescent staining, while nuclear stain DAPI is blue. Scale bars represent 200 microns. (B) Co-immunofluorescence for α-SMA (red) and EP4 (green) demonstrate very few α-SMA^+^, EP4^+^ cells (yellow arrow) in WT repairs at day 21. In contrast, abundant α-SMA^+^, EP4^+^ cells (yellow arrows) are observed in EP4cKO^S100a4^ repairs at this time. Scale bars represent 50 microns. (D-F) Quantitative assessment of (D) α-SMA^+^ cells, (E) EP4^+^ cells, and (F) α-SMA, EP4 double positive cells by immunofluorescence at day 21. (**) Indicates p<0.01 between WT and EP4cKO^S100a4^ repairs.

## Discussion

In the present study we have examined the effects of deleting the PGE2 receptor, *EP4*, specifically in *S100a4*-lineage cells during flexor tendon healing. Given the strong association between inflammation and scar tissue formation ^18^, we tested the hypothesis that loss of *EP4* in *S100a4*-lineage cells would decrease adhesion formation and improve tendon gliding function. Our data demonstrate that EP4cKO^S100a4^ improves tendon gliding at 14 days post-surgery, consistent with a significant decrease in *EP4* expression prior to this point. Conversely, gliding function during the later stages of healing (day 28) is impaired, while *EP4* expression is similarly increased prior to these functional deficits. Taken together, these data suggests that early deletion of *EP4* in S100a4-lineage cells alters the kinetics of healing, potentially resulting in changes in the cellular milieu, and up-regulation of *EP4* by an alternative cell population during later healing. To determine the effects of *EP4* deletion in a different cell population, we examined healing in mice with *EP4* deletion in *Scleraxis-lineage* cells and found no effect on gliding function or mechanical properties. We then examined α-SMA^+^ myofibroblasts as potential cell mediator of increased *EP4* expression and adhesion formation during late healing in EP4cKO^S100a4^ mice. A substantial increase in α-SMA+, EP4+ double-positive cells were observed in EP4cKO^S100a4^, suggesting that an influx of EP4-expressing α-SMA+ myofibroblasts may drive the up-regulation of EP4, potentially re-activating inflammation and promoting scar-mediated tendon healing.

Immediately following tendon injury the acute inflammatory phase begins. While inflammation is necessary for the initiation and progression of the normal healing cascade, there is clear evidence that inflammation can also drive the scar tissue healing response in tendon. Embryonic ^19,20^ and early post-natal^17^ healing are characterized by scarless, regenerative healing in contrast to healing in mature tissues which tend toward a scar tissue response. Changes in the inflammatory and immune environment have been proposed as a main factor in this differential response as the embryonic and early post-natal environment has an altered immune system, relative to adult^21,22^. Furthermore, MRL/MPJ mice, which have a blunted inflammatory response ^23^, also demonstrate improved tendon healing ^24^. Therefore, modulating inflammation represents a promising approach to improve tendon healing. Cox-2 inhibitors have shown promising inhibition of adhesion formation ^2,3^, however systemic Cox-2 inhibitors are plagued by side-effects, including increased risk of adverse cardiovascular events, that must be given serious consideration ^25,26^. Thus, we have focused on downstream targets of Cox-2 mediated inflammation, specifically the Prostaglandin E2 receptor, EP4. We have previously demonstrated that systemic antagonism of EP4 impaired gliding function during tendon healing. However, several studies have demonstrated the differential effects of EP4 in a cell-type dependent context. Yokoyama *et al.,* demonstrated that pharmacological antagonism of EP4, or genetic knock-down of EP4 decreased abdominal aortic aneurysms ^27^. However, Tang *et al.,* demonstrate that bone marrow cell-specific deletion of EP4 promotes abdominal aortic aneurysms ^12^. Taken together these data support investigating cell-type specific effects of EP4 deletion on tendon healing.

The differential effects of systemic versus cell-type specific EP4 inhibition are demonstrated by alterations in ECM gene expression. EP4 antagonist treatment resulted in earlier expression of *Col3a1,* relative to vehicle treated repairs ^10^, suggesting that systemic EP4 antagonism promotes an accelerated shift toward scar-mediated healing. In contrast, *Col3a1* was significantly decreased in EP4cKO^S100a4^ repairs, relative to WT repairs, consistent with improved gliding in EP4cKO^S100a4^ repairs at that time-point. However, expression of *Col1a1,* which is associated with restoration of native tendon composition, is also decreased in EP4cKO^S100a4^ repairs throughout healing, suggesting that EP4cKO^S100a4^ disrupts the normal progression of tendon healing. Based on gliding function, mechanics and gene expression data, WT repairs progress through the normal phases of healing, with progressive acquisition of mechanical properties, a transient decrease in gliding function, and a shift from early *Col3a1* expression to *Col1a1.* In contrast, no change in gliding function is observed over the course of healing in EP4cKO^S100a4^, and diminished acquisition of mechanical properties occurs; max load at failure increases 102% in WT repairs from day 21 to day 28, compared to a 1.6% increase in max load over the same time-period in EP4cKO^S100a4^ repairs. Indeed, following an early failure to form adhesions, EP4cKO^S100a4^ repairs seem to remain relatively static, as no change in GR or MTP Flexion angle is observed over time, and no up-regulation of *Col3a1* or *Col1a1* is observed. Taken together, these data may suggest that early deletion of *EP4* in S100a4-lineage cells halts progression through the normal phases of tendon healing.

Given the differences in healing between EP4 antagonist treated mice, and EP4cKO^S100a4^ mice, we examined the effects of EP4 conditional deletion in Scleraxis-lineage cells. *Scleraxis* is required for normal tendon formation^28^, and is expressed by resident tenocytes in mature tendon^17^. Recently, *Scx*-lineage cells have been shown to participate in regenerative tendon healing in neonates, with minimal involvement of *Scx^+^*cells in mature, scar mediated healing ^17^, which may account for the lack of phenotype observed in EP4cKO^Scx^. Furthermore, while *Scx*-lineage cells are observed in the healing tendon (Figure 6), we have used a noninducible *Scx*-Cre, so it is unknown if these *Scx*-lineage^+^ cells are residual cells from development, or newly activated *Scx*-expressing cells that are involved in healing.

Myofibroblasts are believed to be the cellular drivers of fibrotic tendon healing and peritendinous adhesions ^29,30^, given their ability to produce abundant collagenous ECM. We have used α-SMA expression to identify myofibroblasts, consistent with many other studies ^31–33^. However, α-SMA has also been used to mark mesenchymal progenitors ^34^. While future studies will clarify the lineage, fate and function of these cells during tendon healing, there is a consistent demonstration of the involvement of α-SMA+ cells during tendon healing. While our data suggest that α-SMA+ myofibroblasts drive the up-regulation of EP4 that occurs during later healing in EP4cKO^S100a4^, it is not clear if the increase is driven by elevated levels of the EP4 ligand PGE2, and which cells are producing PGE2 at this stage in healing. Macrophages play important roles in both initiation and resolution of inflammation, and have been shown to produce PGE2 ^35^. However, PGE2 can also alter macrophage polarization ^36,37^, and recent work by Fernando *et al.,* demonstrate that myofibroblast-derived PGE2 can promote alternative activation of macrophages ^38^. Thus, understanding the relationship between myofibroblasts, macrophages and PGE2/ EP4 signaling during tendon healing will be an important area of future research.

While these data clearly identify different cell-type specific functions of EP4 in tendon healing there are several limitations that must be considered. We have examined the effects of EP4cKO^Scx^ at only one time-point during healing, however, if EP4cKO^Scx^ were to alter healing, our previous data would suggest these effects would occur prior to day 14 based on the expression profile of *Scx* in this healing model ^39^. We have also not examined the long-term mechanical effects of EP4cKO^S100a4^. on tendon healing. In addition, the actual identity of S100a4+ cells is both unclear and controversial. Osterreicher *et al.,* have identified S100a4+ cells as a subpopulation of inflammatory macrophages ^40^, while Nishitani *et al.,* identified S100a4+ cells as myofibroblasts ^41^. Thus, future studies to more clearly define the origin, fate and function of S100a4 during tendon healing are needed. In addition, we have not examined the role of other EP receptors in EP4cKO^S100a4^ repairs, which will be an important consideration in future studies, given that loss of EP2 is associated with fibrotic progression in the lung ability of EP2 in particular to also promote fibrosis ^42,43^. Finally, while these data suggest a role for α-SMA^+^ cell-mediated EP4 induction, and increased scar formation, conditional deletion studies of EP4 using an α-SMA-Cre mouse would be required to definitively identify the role of *EP4* in *α-SMA-lineage* cells.

Together with our previous studies using an EP4 systemic antagonist, these data demonstrate cell-type specific functions of EP4 in scar-mediated tendon healing. Our data suggest that deletion of EP4 in *S100a4-*lineage cells inhibits range of motion-limiting scar tissue without further compromising mechanical properties. However, these beneficial effects of EP4cKO^S100a4^ are transient, with aberrant up-regulation of EP4 during later healing, mediated at least in part by α-SMA+ cells, and a subsequent decrease in range of motion. These data suggest that sustained inhibition of EP4 in S100a4-lineage cells, and *α-SMA-lineage* cells may result in prolonged improvements in tendon healing. Furthermore, understanding the mechanisms through which EP4 promotes scar-mediated healing will identify novel therapeutic targets to improve tendon healing.

## Methods

### Animal Ethics

This study was carried out in strict accordance with the recommendations in the Guide for the Care and Use of Laboratory Animals of the National Institutes of Health. All animal procedures were approved by the University Committee on Animal Research (UCAR) at the University of Rochester (UCAR Number: 2014-004).

### Mice

The following strains were obtained from Jackson Laboratories (Bar Harbor, ME): *EP4*^flox/flox^ (#28102), *ROSA*^nT/nG^ (# 23035) *s100a4-Cre* (#12641). The *S100a4-Cre* ^44^ mice were generated on a BALB/cByJ background and were back-crossed to C56Bl/6J for at least 10 generations prior to crossing to *EP4*^flox/flox^ or *ROSA*^nT/nG^ strains. *Scx*-Cre mice were generously provided by Dr. Ronen Schweitzer. Conditional knock-out animals were generated as homozygotes (Cre^+^; *EP4*^flox/flox^), Cre^-^; *EP4*^flox/flox^ littermates were used as wildtype (WT). The *ROSA*^nT/nG^ construct is a dual reporter in which all cells express nuclear Tomato red (nT) fluorescence in the absence of Cre-mediated recombination, and nuclear GFP (nG) upon Cre-mediated recombination.

### Flexor Tendon Surgery

At 10-12 weeks of age mice underwent complete transection and repair of the flexor digitorum longus (FDL) tendon in the right hind-paw as previously described ^39,45^. Briefly, mice were anesthetized with Ketamine (60mg/kg) and Xylazine (4mg/kg). A pre-surgical dose of Buprenorphine (0.05mg/kg) was administered followed by further analgesia every 12-hours after surgery as needed. Following preparation of the surgical site, the FDL was surgically transected in the transverse plane at the myotendinous junction in the calf to protect the repair site from high strains. The skin was closed with a single 5-0 suture. A 1-2cm incision was then made on the posterior surface of the hindpaw, soft tissue was retracted to identify the FDL, and the FDL was completely transected using micro-scissors. Following transection the FDL was repaired using 8-0 nylon sutures (Ethicon, Somerville, NJ) in a modified Kessler pattern. The skin was closed with 5-0 suture. The animals were allowed unrestricted weight-bearing and movement, and had *ad-libitum* access to food and water.

### RNA extraction and qPCR

Total RNA was extracted from healing tendons between 3-28 days post-surgery. RNA was extracted from the repair site and 1-2mm of native tendon on both sides of the repair. RNA was extracted from 3 repairs per genotype per time-point. RNA was extracted using TRIzol reagent (Life Technologies). 500ng of RNA was used for reverse transcription of cDNA using iScript cDNA synthesis kit (Bio-Rad, Hercules, CA). Quantitative PCR (qPCR) was then run using cDNA and gene specific primers (Table 1). Expression was normalized to expression in day 3 WT repairs, and the internal control *β-actin.* All qPCR was run in triplicate and repeated at least twice.

**Table 1.**
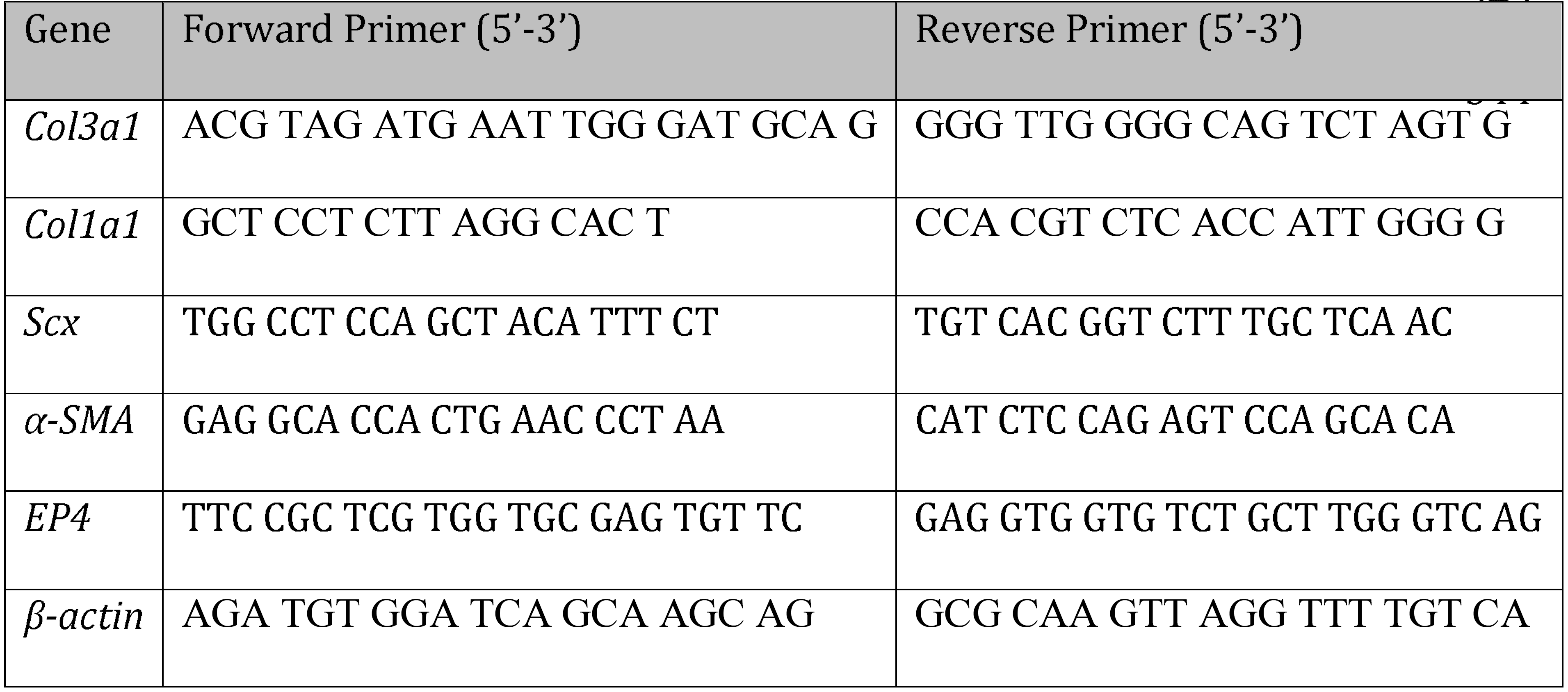
Primer Sequences.

### Histology-Frozen Sectioning

Following sacrifice the hind limbs of *S100a4*-Cre^+^; *ROSA*^nT/nG^ and *Scx*-Cre; *ROSA*^nT/nG^ mice were removed at the mid-tibia, and skin and excess soft tissue was removed. The skin on the sole of the foot remained intact so as not to disturb the repair site. Tissues were fixed in 10% neutral buffered formalin for 24 hrs at 4°C, decalcified in 14% EDTA (pH 7.2-7.4) for 4 days at 4°C, and processed in 30% sucrose (in PBS) for 24 hours prior to embedding in Cryomatrix (ThermoFisher, Waltham, MA). Eight-micron serial sagittal sections were cut using a cryotape-transfer method ^46^. Sections were affixed to glass slides using 1% chitosan in 0.25% acetic acid, dried overnight at 4°C, and coverslipped using Fluoromount aqueous mounting medium (F46802, Sigma, St. Louis, MO. Following fluorescent imaging, coverslips were gently removed in PBS, and slides were stained with Hematoxylin & Eosin, followed by dehydration and coverslipping. Both fluorescent and brightfield imaging was done on a VS120 Virtual Slide Microscope (Olympus, Waltham, MA). Images are representative of at least three specimens per time-point.

### Histology-Paraffin sectioning

Hind limbs were removed as above and fixed for 72hrs in 10% NBF at room temperature. Specimens were then decalcified for 14 days in 14% EDTA (pH 7.2-7.4), processed and embedded in paraffin. Three-micron serial sagittal sections were cut, de-waxed, rehydrated, and probed with antibodies for EP4 (1:100, #sc-16022, Santa Cruz Biotechnology, Dallas, TX) and α-SMA-Cy3 (1:200, #C6198, Sigma, St. Louis, MO) overnight at 4°C. An AlexaFlour 488 secondary antibody was used with EP4 (1:200, #705-546-147, Jackson ImmunoResearch, West Grove, PA). The nuclear counterstain DAPI was used, and fluorescence was imaged with a VS120 Virtual Slide Microscope (Olympus, Waltham, MA). Following imaging, coverslips were removed by immersing slides in PBS, and then slides were stained with Hematoxylin & Eosin (H&E).

### Quantification of fluorescence

Digital fluorescent images obtained from the slide scanner were processed using Visiopharm image analysis software v.6.7.0.2590 (Visiopharm, Hørsholm, Denmark) Automatic segmentation was done using the threshold classifier to define cell populations based on fluorescent channel. A double constraint was defined to determine co-localization of EP4 and α-SMA staining. Regions of interest were drawn to include only the scar tissue area for processing. To improve accuracy of quantification, manual correction was done to exclude sutures/blood vessels, and a post-processing step was added to ignore any debris smaller than 10μm^2^. The area of each fluorescent signal was calculated, and these values were used to determine overall percentage of each cell type or stain present in the scar tissue.

### Assessment of Gliding Function

Following sacrifice the hindlimb was removed at the knee, and skin was removed down to the ankle. The FDL was isolated proximally near the original myotendinous junction and secured using cyanoacrylate between two pieces of tape as previously described ^39,47^. Briefly, the tibia was secured in an alligator clip and the tendon was incrementally loaded with small weights from 0-19g; digital images were taken at each load. The flexion angle of the metatarsophalangeal (MTP) joint was measured from the digital images. The MTP Flexion angle corresponds to the degrees of flexion upon application of the 19g weight. Un-injured tendons undergo complete flexion of the digits at 19g. Gliding resistance was based on the changes in MTP flexion angle of the range of applied loads with higher gliding resistance indicative of impaired gliding function and adhesion formation. Eight-10 specimens per genotype per time-point were analyzed for gliding function.

### Biomechanical testing

Following Gliding testing, the tibia was removed at the ankle and the toes and proximal section of the FDL in the tape were secured in opposing ends of a custom grips ^39,47^ on an Instron 8841 DynaMight^TM^ axial servohydraulic testing system (Instron Corporation, Norwood, MA). The tendons were tested in tension at a rate of 30mm/minute until failure. Maximum load at failure was automatically logged, while stiffness was calculated at the slope of the linear region of the force-displacement curve. Eight-10 specimens per genotype per time-point underwent mechanical testing.

### Statistical analyses

Quantitative data are presented at mean ± standard error of the mean (SEM). A two-way analysis of variance (ANOVA) with Bonferroni’s multiple comparisons test was used to analyze qPCR, biomechanical and gliding data of WT and EP4cKO^S100a4^ tendon repairs over time. A one-way ANOVA with Bonferroni’s multiple comparisons test was used to analyze nGFP+ quantification. A t-test was used to analyze biomechanical and gliding data from un-injured tendons, EP4cKO^Scx^ vs. WT repairs at day 14, and quantification of immunofluorescence.

## Supplemental Figures

**Supplemental Figure 1.** Hematoxylin & Eosin (H & E) staining of (A) un-injured, and repaired tendons at (B) 7, and (C) 14 days post-surgery from *S100a4*-Cre+; nT/nG mice. Following fluorescent imaging (Figure 1), coverslips were gently removed and sections were stained with H&E to assess tissue and cell morphology. Tendon is outlined in black. Scale bars represent (A) 100 microns, (B&C) 200 microns.

**Supplemental Figure 2.** Hematoxylin & Eosin (H & E) staining of (A) un-injured tendon and (B) day 14 postsurgery from Scx-Cre+; nT/nG mice. Following fluorescent imaging (Figure 6), coverslips were gently removed and sections were stained with H&E to assess tissue and cell morphology. Tendon is outlined in black. Scale bars represent 200 microns.

**Supplemental Figure 3.** Hematoxylin & Eosin (H & E) staining of (A & A’) WT and (B & B’) EP4cKO^S100a4^ tendon repairs at day 21 post-surgery. Following fluorescent imaging (Figure 7), coverslips were gently removed and sections were stained with H&E to assess tissue and cell morphology. Tendon is outlined in black. Scale bars represent 200 microns (A & B), or 50 microns (A’ & B’). Blood vessels are identified by black arrows.

**Supplemental Figure 4.** To assess changes in α-SMA and EP4 protein expression through the depth of the repair site between WT and EP4cKO^S100a4^ repairs, four sections that were approximately 40 microns apart (spanning 120 microns of healing tissue) underwent co-immunoflourescent staining for α-SMA (red) and EP4 (green). Nuclei are stained blue with DAPI. Scale bars represent 50 microns. Red arrows indicate blood vessels in the H & E images.

## Acknowledgements

We would like to thank the Histology, Biochemistry and Molecular Biology (HBMI) and Biomechanics Multimodal Tissue Engineering (BMTI) Cores in the Center for Musculoskeletal Research (CMSR) at the University of Rochester for technical assistance with histology and biomechanical testing, respectively.

This work was supported in part by NIH/ NIAMS 1K01AR068386-01 (to AEL), and R01 AR056696. The HBMI and BMTI Cores were supported in part by NIH/ NIAMS P30 AR069655.

## Author contributions

Study conception and design: RJO, AEL; Acquisition of data: JEA, KTB; Analysis and interpretation of data: JEA, KTB, AEL; Drafting of manuscript: JEA, AEL; Revision and approval of manuscript: JEA, KTB, RJO, AEL.

## Data Availability Statement

All data generated or analysed during this study are included in this article.

